# JNK and PI3K signaling pathways mediate synapse formation and network spontaneous activities in primary neurons

**DOI:** 10.1101/2024.04.23.590713

**Authors:** Xiaoli Jia, Qiuyan Zhu, Shaohua Wu, Zhihong Zhou, Xian Jiang

**Affiliations:** Institute of Neurological and Psychiatric Disorders, Shenzhen Bay Laboratory, Shenzhen, 518000, China; School of Chemical Biology and Biotechnology, Peking University, Shenzhen, 518055, China; School of Medicine, The Chinese University of Hong Kong, Shenzhen, Shenzhen, Guangdong, 518172, China

**Author notes:** Corresponding authors.; E-mail addresses. J.X.L, Z.Q.Y and W.S.H. contributed equally to this study.

**Keywords:** Synapse formation, C-jun N-terminal kinases, Phosphatidylinositide3-kinases, Neural network function, Neural network development

## Abstract

**Background:** Cellular signals orchestrating synapse formation and neuronal network function remain poorly understood. To explore the critical signaling pathways in neurons and their influence on network development, pharmacological assays were employed to inhibit multiple signaling pathways in cultured neurons.

**Methods:** Immunofluorescence and western blotting are applied to identify the expression of synapse-related proteins within neurons. micro-electrode arrays (MEAs) are employed to study the developmental characteristics of neuronal networks. RNA sequencing is utilized to determine the gene expression profiles pertaining to multiple signaling pathways.

**Results:** Canonical c-jun N-terminal kinases (JNK) pathway is necessary for pre- and post-synaptic specializations, while phosphatidylinositide3-kinases (PI3K) is a key to postsynaptic specialization and affects the puncta sizes of presynaptic marker. Unexpectedly, pharmacological inhibition of JNK pathway significantly suppressed the mean firing rate (MFR), network burst frequency (NBF) and regularity of network firing after 4 weeks, but did not alter the synchrony of the network. During network development, PI3K pathway regulates the longer burst duration and lower network synchrony. Gene sets associated with neurodevelopmental processes and myelination was disturbed during restraining these signal pathways. Furthermore, inhibition of the PI3K signaling pathway obviously transformed voltage-gated ion channel activity, synaptic transmission and synaptic plasticity of neurons.

**Conclusion:** This study reveals that JNK and PI3K signaling pathways play different roles during synapse formation, and these signaling pathways have a lasting impact on the development of neuronal networks. Thus, this study provides further insights into the intracellular signaling pathways associated with synapse formation in the development of neuronal networks.

## Introduction

Despite synapse formation being a universal and essential process for constructing all brain circuits, its mechanism of action (MOA) remains unclear, including the intracellular signaling pathways involved in assembly of pre-/post-synaptic specializations, the potential existence of different signaling pathways for these specializations, and the organization of excitatory and inhibitory synapses. Synapses exhibit the classic characteristic that neurotransmitters are rapidly and transiently released from the presynaptic side and recognized by the postsynaptic side (*1*). It’s intriguing that canonical features shared by all synapses, like synaptic vesicles and active zones, are exclusive to the presynaptic side. These components are identical in both excitatory and inhibitory synapses. However, the postsynaptic sides of excitatory synapses distinctly differed from inhibitory synapses (*2*). Even excitatory and inhibitory neurotransmitter receptors lack homology, sharing few molecular components between their respective postsynaptic specializations. In artificial synapse formation assays which offer a viable method to survey the synapse formation involved signaling pathways (*3*) (*4*), synaptic adhesion molecules transfected in a HEK93 or non-neuronal cell drive the pre- or post-synaptic specializations in co-cultured neurons (*5*).

Pharmacological inhibitors and genetic tools aid in elucidating the contributions of c-jun N-terminal kinases (JNK) and protein kinase-A (PKA) signaling to the formation of both pre- and post-synaptic specializations in heterologous synapse formation (HSF). Conversely, the phosphatidylinositide3-kinases (PI3K) signaling pathway is specifically required for the formation of postsynaptic specializations without concurrent presynaptic specializations (*4*). Furthermore, *in vitro* neuronal models have emerged as crucial tools for investigating the intricate communication between different neuronal conditions. Specifically, the capacity to observe and manipulate the firing activity of neuronal network offers valuable insights into neuronal population development and organization (*6*). Under this circumstances, micro-electrode arrays (MEAs) were utilized to record spontaneous neuronal network activity, aiming at exploring the involvement of various signaling pathways in neuronal network development and synapse formation. MEAs, cell culture dishes equipped with embedded micro-electrodes, facilitates noninvasive measurements of neuronal network activity (*7*) (*8*) (*9*), and their effectiveness in measuring the response of primary neurons to medicines has been demonstrated previously (*10*) (*11*) (*12*). However, little is known about how intracellular JNK, PKA and PI3K/AKT signaling pathways affect the firing activity and development of neuronal network.

In the present study, pharmacological inhibitors and electrophysiological approaches were applied to explore the internality of the signals mediates endogenous synapse formation and neuronal function. Multiple signaling pathways play an important role in the pre- and post-synaptic specializations in culture neurons, especially the mean firing rate (MFR) in JNK of the neuronal network, network bursts (NBs) and NBs duration, network synchrony of PI3K. Inhibition of the JNK and PI3K pathways altered transcriptomic profile. Thus, an initial insight was present into the signaling mechanisms underlying cultured neuron electrophysiology that involve in synapse formation and neuronal function.

## Materials and Methods

### Chemicals and pharmacological agents

The following compounds were used: H-89 dihydrochloride (Med; Cat# 127243850, 1 μM); PKI (Millipore; Cat# 476485, 10 μM); JNK-IN-8 (Selleck; Cat# S4901, 0.3 μM); SP6000125 (Sigma; Cat#S5567, 1 μM); GDC-0941(Apexbio; Cat# A8210, 0.3 μM); LY294002 (Sigma; Cat# L9908, 3 μM).

### Antibodies

The subsequent antibodies were employed at the specified concentrations: anti-MAP2 Guinea pig (Synaptic Systems; Cat# 188004; 1:1000); anti-VGAT mouse (Synaptic Systems; Cat# 131011; 1:1000); anti-VGluT1 rabbit (Synaptic Systems; Cat# 135316; 1:1000); anti-PSD95 mouse (Thermo Fisher; Cat# MA1046; 1:500); phospho-CREB (Cell Signaling Technology; Cat# 9198S; 1:1000); phospho-JNK (Cell Signaling Technology; Cat# 9255S; 1:2000); phospho-Akt (Cell Signaling Technology; Cat# 1911; 1:2000); Goat anti-Guinea Pig Secondary Antibody, Alexa Fluor™ 488 (Thermo Fisher; Cat# A11037; 1:1000); Goat anti-Rabbit Secondary Antibody, Alexa Fluor™ Plus 647 (Thermo Fisher; Cat# A32733; 1:1000); Goat anti-Mouse Secondary Antibody, Alexa Fluor™ 546 (Thermo Fisher; Cat# A11030; 1:1000).

### Primary neurons cultures

Similar to previous descriptions (*13*), rat cortical neurons were prepared, with careful exclusion of the hippocampus from the dissected cortex. Briefly, embryos were aseptically removed and dissected. The cortex was dissociated through enzymatic digestion in 0.125% trypsin (Thermo Fisher, United States) for 10 minutes at 37 ^◦^C, followed by trituration with a Pasteur pipette. The dissociated neurons were then plated onto Poly-D-lysine-coated glass coverslips in 24-well plates. Four hours later, after cell adherence to the substrate, 1 mL of medium was added to each well. The cells were cultured in neurobasal plus medium (Invitrogen, United States) supplemented with 1% GlutaMAX (Gibco, United States) and 2% B-27 plus (Gibco, United States) in a humidified atmosphere of 5% CO at 37 ^◦^C. Starting from DIV7, 30% of the medium was replaced every 3 days until analysis at DIV14-DIV16.

### Immunocytochemistry

All immunocytochemistry experiments were conducted as previously described (*14*). Neurons were briefly washed once with PBS, fixed with 4% PFA for 20 minutes at room temperature (RT), and then washed three times for 10 minutes each in PBS. Subsequently, they were permeabilized in 0.2% Triton 100/PBS for 10 minutes at RT. Following this, cells were blocked in blocking buffer (5% BSA/PBS) for 30 minutes and incubated overnight at 4 °C with diluted primary antibodies in blocking buffer. After the primary antibody incubation, cells were washed three times for 10 minutes each in PBS, then incubated with secondary antibodies in blocking buffer for 2 hours. Finally, they were washed three times in PBS and mounted on microscope slides using Fluoromount-G (Southern Biotech).

### Image acquisition and analysis

Neuronal morphology images were captured using a ZEISS LSM980 confocal microscope, maintaining constant laser gain and offset settings, scanning speed, and pinhole size (*15*). These settings ensured that images did not have oversaturated pixels. Each z stack consisted of 10 serial images acquired in 0.5 mm steps, and maximum intensity projections of the z stacks were generated for quantification. Synaptic puncta were counted along well-isolated primary dendrites using IMARIS (OXFORD Instruments). To measure dendritic length and branching, field images of low-density neuronal cultures, counterstained with anti-MAP2, were analyzed using IMARIS. Constant threshold settings were maintained across all experimental conditions to exclude background signals.

### Immunoblotting

Cells and neurons were lysed in RIPA buffer (50 mM Tris-HCl pH 8.0, 150 mM NaCl, 0.1% SDS, 0.5% sodium deoxycholate, 1% Triton X-100), supplemented with a mixture of protease inhibitors (Roche) (*16*). Lysates were subjected to SDS-PAGE in the presence of dithiothreitol (DTT, 0.1 M). Immunoblotting and quantitative analysis were conducted using fluorescent-labeled secondary antibodies and an Odyssey Infrared Imager CLX with Image Studio 5.2.5 software (LI-COR Biosciences). Signal normalization was performed using neuronal GAPDH probed on the same blots as loading controls.

### MEA recording

Using the 6-well MEA system (AXION Biosystems, Maestro Pro, America), all recordings were conducted as previously described (*17*). These MEA devices facilitate non-invasive recording of neuronal activity simultaneously at multiple nodes within the same network. Each of the 6 independent wells is equipped with 64 micro-electrodes. Neuronal networks were allowed to acclimate to the recording environment (37◦C; 5% CO_2_) for 10 minutes, following which spontaneous neuronal network activity was recorded for an additional 10 minutes. The signal was sampled at 10 KHz, filtered with a high-pass filter (Butterworth, 100 Hz cut-off frequency), and the noise threshold was set at ± 5 standard deviations.

Offline electrophysiological data analysis was conducted using AxlS Navigator software (AXION Biosystems), allowing for the extraction of spike-trains, bursts, NBs, and other electrophysiology parameters. Batch processing was carried out using custom in-house code developed in MATLAB (MathWorks, Natick, MA, USA). To ensure data quality, the following criteria were applied for exclusion from analysis: 1) neuronal network activity with a mean firing rate (MFR) < 0.1 Hz and burst rate (BR) < 0.4 bursts/min, 2) cultures lacking NBs at DIV 28, and 3) low cell density or cell clustering.

### RNA-Seq

Total RNA was extracted using the Trizol reagent kit (Invitrogen, Carlsbad, CA, USA) following the manufacturer’s protocol. RNA quality was evaluated on an Agilent 2100 Bioanalyzer (Agilent Technologies, Palo Alto, CA, USA) and checked via RNase-free agarose gel electrophoresis. Following total RNA extraction, eukaryotic mRNA was enriched using Oligo(dT) beads. The enriched mRNA was fragmented into short fragments using fragmentation buffer and reverse transcribed into cDNA using the NEBNext Ultra RNA Library Prep Kit for Illumina (NEB #7530, New England Biolabs, Ipswich, MA, USA). The resulting double-stranded cDNA fragments were subjected to end repair, addition of A bases, and ligation to Illumina sequencing adapters. The ligation reaction was purified using AMPure XP Beads (1.0X). Ligated fragments underwent size selection via agarose gel electrophoresis and were amplified by polymerase chain reaction (PCR). The resulting cDNA library was sequenced using the Illumina Novaseq6000 by Gene Denovo Biotechnology Co. (Guangzhou, China).

### Statistical analysis

Using Prism 8 software (GraphPad Software), statistical analyses were conducted. Quantitative data presented in the figures are expressed as mean ± SD. Each experiment was repeated independently at least three times. Unpaired *t*-tests or one-way ANOVAs with Tukey’s post hoc tests were employed for all statistical analyses, comparing control and treated conditions within the same experiments.

## Results

### JNK and PI3K signal pathway did not inhibited neuronal morphology

The cortical neurons were extracted from 18th day embryonic mice and related intracellular signaling pathways were pharmacologically inhibited at 7-14 days (Figure 1A). Gradient screening was employed to determine the minimum inhibitory concentrations (MICs) for each inhibitor. On the 7th day of neuron culture, inhibitors were added basing on their MICs, and neuronal proteins were extracted after one week of treatment. The MICs of PKA inhibitors H89 and PKI, JNK inhibitors JNK-IN-8 and SP6000125, PI3K inhibitors GDC-0941 and LY294002 were 1 μM, 10 μM, 0.3 μM, 10 μM, 0.3 μM and 3 μM, respectively (Figure S1). Immunofluorescence of the DIV 15 neurons showed that PKA inhibitors H89 and PKI significantly decreased the size of soma, the number of primary dendrites and total neurite length (Figure 1B, F-H, *P*<0.05). However, the size of the neuronal soma, the number of primary dendrites and neuron length did not change markedly in the JNK and PI3K pathway (Figure C-H, *P*>0.05). Excessive changes in the growth and development of neurons irreversibly disrupt the signaling pathway and affect the maintenance of the whole cell homeostasis. Thus, only JNK and PI3K pathway were used to study synapse formation in further experiments, in order to avoid such factors interfering the accuracy of experimental results.

**Figure 1.**
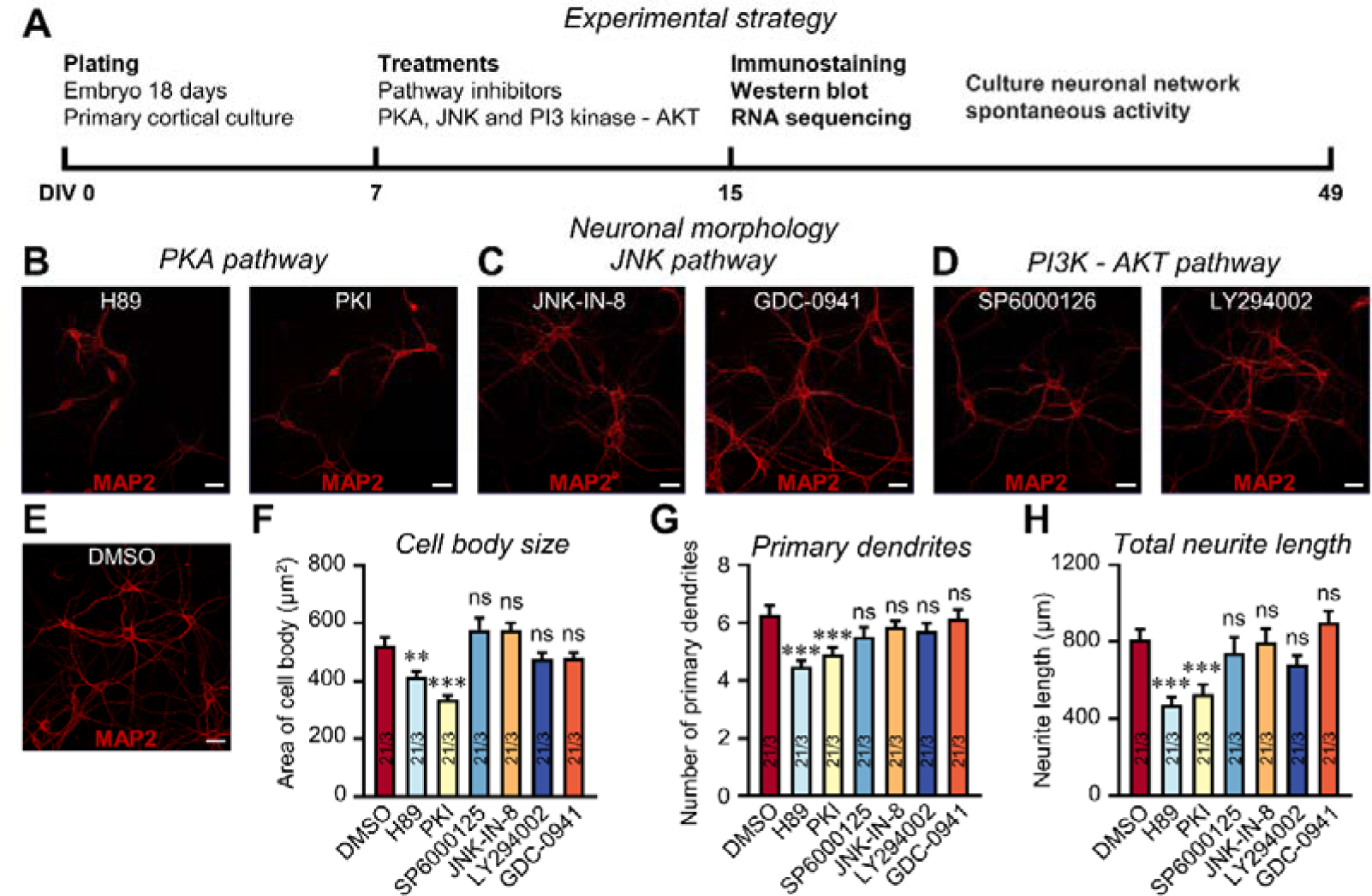
JNK and PI3K signaling pathway show no effect on morphology in primary cultures of cortical neurons. **A**. Experiment strategy. Cortical neurons were cultured from E18 wide type mice. At DIV7, neurons were treated with PKA, JNK and PI3K pathway inhibitors. Then, neurons were analyzed by immunocytochemistry, immunoblotting and RNA-seq at DIV15. Neuronal network spontaneous activities were recorded from DIV15 to DIV28. **B**. PKA inhibitors (H89, 1 μM; PKI, 10 μM) alter the dendritic morphology of neurons. **C**. JNK inhibitors (SP6000125, 10 μM; JNK-IN-8, 0.3 μM) unchange the dendritic morphology of neurons. **D**. PI3K-AKT inhibitors (LY294002, 3μM; GDC-0941, 0.3μM) do not alter the dendritic morphology of neurons. **E**. DMSO as the control. Neurons were stained with MAP2. Quantifications showing that PKA pathway alter the size of soma (**F**), number of primary dendrites (**G**) and total neuron length (**H**), but JNK and PI3K pathways show no effect on neuronal morphology. All numerical data are means ± SD. Numbers in bars list the number of independent experiments/cells analyzed. Statistical significance was examined by Student’s *t* test (\**P*<0.05, \*\**P*<0.01, \*\*\**P*<0.001).

### JNK pathway interfere pre-/post-synaptic specialization, and PI3K signal only perturb post-synaptic puncta density in the cultured neurons

The immunofluorescence was performed on pre- and post-synaptic markers to investigate potential intracellular signaling pathways governing synapse formation between neurons (Figure 2A). Inhibiting JNK pathway observably decreased synapse density and puncta size (Figure 2B and 2F, *P*<0.05). The PI3K inhibitor LY294002 had no effect on synapse density and size. However, GDC-0941, another PI3K inhibitor, caused an obvious decrease in synapse density and size (Figure 2A, 2B and 2F, *P*<0.05). Furthermore, the roles of the JNK pathway induced presynaptic VGluT1 and VGAT were verified by SP6000125 and JNK-IN-8 (Figure 2C and 2D). Presynaptic VGluT1 and VGAT were severally utilized as substitutes for excitatory and inhibitory synapses. Interestingly, only the JNK antagonist SP6000125 notably lessened the size of VGluT1 puncta (Figure 2G and 2H, *P*<0.05). JNK blocker SP6000125 and JNK-IN-8 also signally suppressed the post-synaptic specializations for the decrease of PSD95 puncta density and size (Figure 2E and 2I, *P*<0.05). Neither the densities of excitatory nor inhibitor pre-synaptic specialization was impressed by PI3K inhibitor (Figure 2C and 2D, *P*>0.05). Only the PI3K inhibitor LY294002 did not significantly reduce the size of the VGAT (Figure 2G and 2H, *P*>0.05). In contrast, PI3K inhibitors LY294002 and GDC-0941 observably intercepted post-synaptic specialization (Figure 2E and 2I, *P*<0.05). The canonical PKA pathway was identified as essential components for assembly of both pre- and post-synaptic specializations (Figure S2). Either pre- or post-synaptic specialization depended on JNK and PKA signals, but the PI3K pathway is merely particularly important for post-synaptic specialization (Figure 2 L-P).

**Figure 2.**
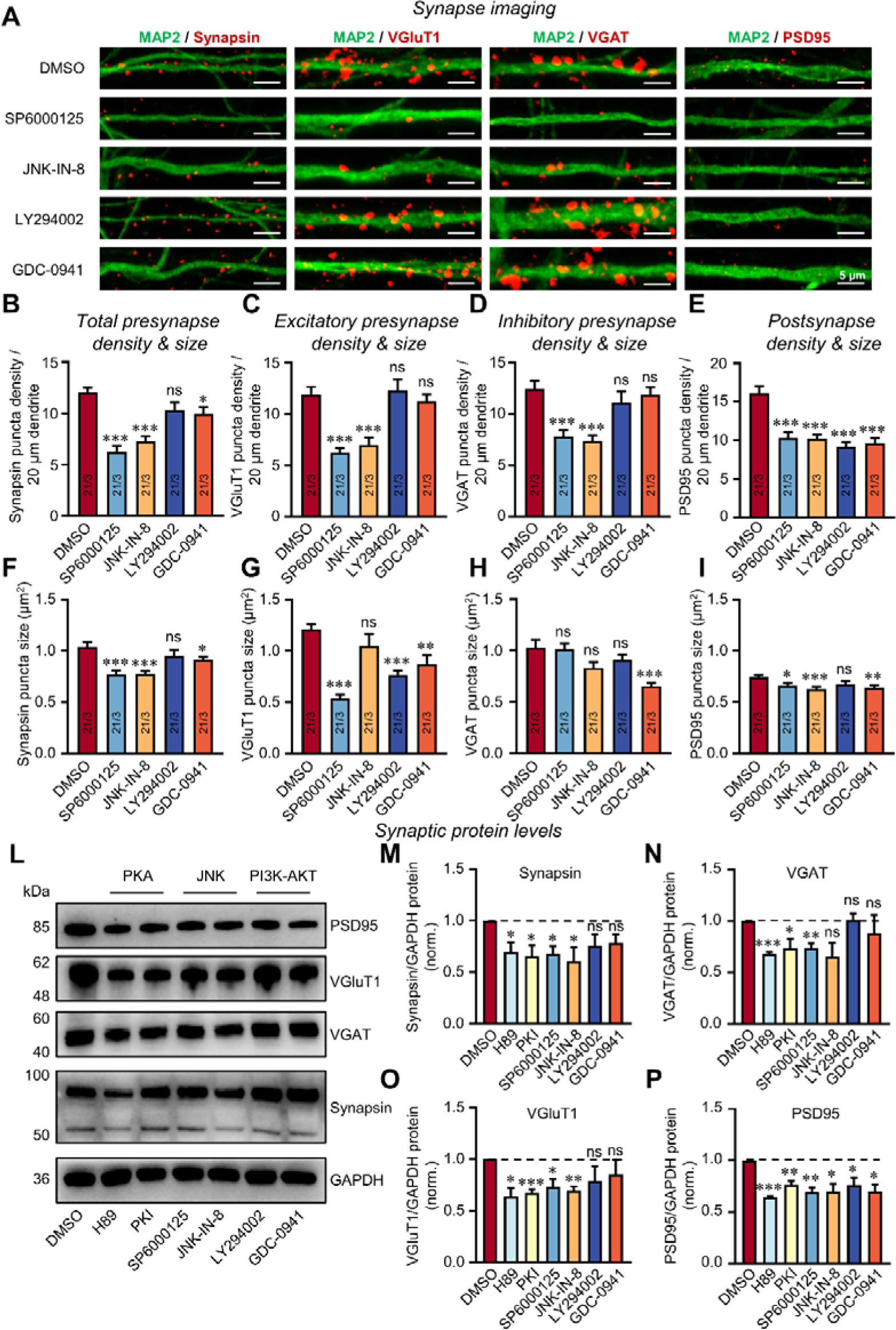
JNK inhibitors impair formation of pre- and post-synaptic sides, while PI3K inhibitors suppress formation postsynaptic density in primary cultures of cortical neurons. **A**. JNK inhibitors treatment decrease both pre- and post-synaptic puncta density, while PI3K inhibitors decrease post-synapse specialization. Neurons were immunostaining for Synapsin1/2 (synapse marker), VGluT1 (excitatory presynaptic marker), VGAT (inhibitory presynaptic marker), PSD95 (postsynaptic marker) and MAP2 (dendritic marker) at DIV15. **B, F**. JNK inhibitors SP6000125 and JNK-IN-8 decrease total pre-synapse density and size. PI3K inhibitors LY294002 do not alter total pre-synaptic puncta density and size, while another PI3K inhibitor GDC-0941 had significant effect on total pre-synapse density and size. **C, G**. JNK inhibitors decrease excitatory pre-synapse density, but JNK-IN-8 display no effect on excitatory pre-synapse size. PI3K inhibitors do not alter excitatory pre-synapse number, but significant effect on excitatory pre-synapse size. **D, H**. JNK inhibitors decrease inhibitory pre-synapse density but does not alter inhibitory pre-synaspse size. PI3K inhibitors do not influence inhibitory pre-synapse density, but GDC-0941 has significant effect on inhibitory pre-synapse size. **E, I**. JNK and PI3K inhibitors has significant effect on post-synapse density, while LY294002 does not alter post-synapse size. **L**. Synapse markers for PKA, JNK and PI3K were assayed by western blot. **M**. Immunoblotting for synapse of PKA, JNK and PI3K was assayed by western blot with detected bands from 50 to 100 kDa. **N**. Immunoblotting for VGAT of PKA, JNK and PI3K was assayed by western blot with detected bands from 40 to 60 kDa. **O**. Immunoblotting for VGluT1 of PKA, JNK and PI3K was assayed by western blot with detected bands from 48 to 62 kDa. **P**. Immunoblotting for PSD95 of PKA, JNK and PI3K was assayed by western blot with detected bands at 85 kDa. Summary graphs of protein levels normalized for GAPDH as an internal standard. All numerical data are means ± SD. Numbers in bars list the number of independent experiments/cells analyzed. Statistical significance was examined by Student’s *t* test (\**P*<0.05, ** *P*<0.01, \*\*\**P*<0.001, *n*≥3).

### Inhibition of the JNK and PI3K pathways impairs neuronal network activity and development

MEAs non-invasively and continuously record neuronal population activity, including spikes and bursts, through extracellular electrodes positioned at spatially distinct locations throughout primary cultures (Figure 3A). Cultured neuronal networks gradually mature from DIV 21 *in vitro*, at which the neuronal network exhibits steady and regular synchronous NBs activities. These NBs are essential not only for network communication, but also for the growth and maturation of the neural network. To inquiry the impact of different signaling pathways on the maturity of the neuronal network and neuronal activity, the electrophysiological signals of DIV 21 were recorded by MEAs. Suppression of JNK and PI3K signal increased the network MFR (Figure 3B-E, *P*<0.05) and network burst frequency (NBF, Figure 3F, *P*<0.05). JNK pathway inhibitors SP6000125 and JNK-IN-8 markedly decreased network burst duration (NBD, Figure 3C and G, *P*<0.05). Moreover, the inhibition of the PI3K pathway did not affect NBD (Figure 3D and G, *P*>0.05). Electrophysiology showed that inhibitors of JNK and PI3K also had a noteworthy implication on the number of spikes per NBs (Figure 3H). Therefore, the maturation of neuronal networks and the firing patterns was adjusted by JNK and PI3K pathway.

**Figure 3.**
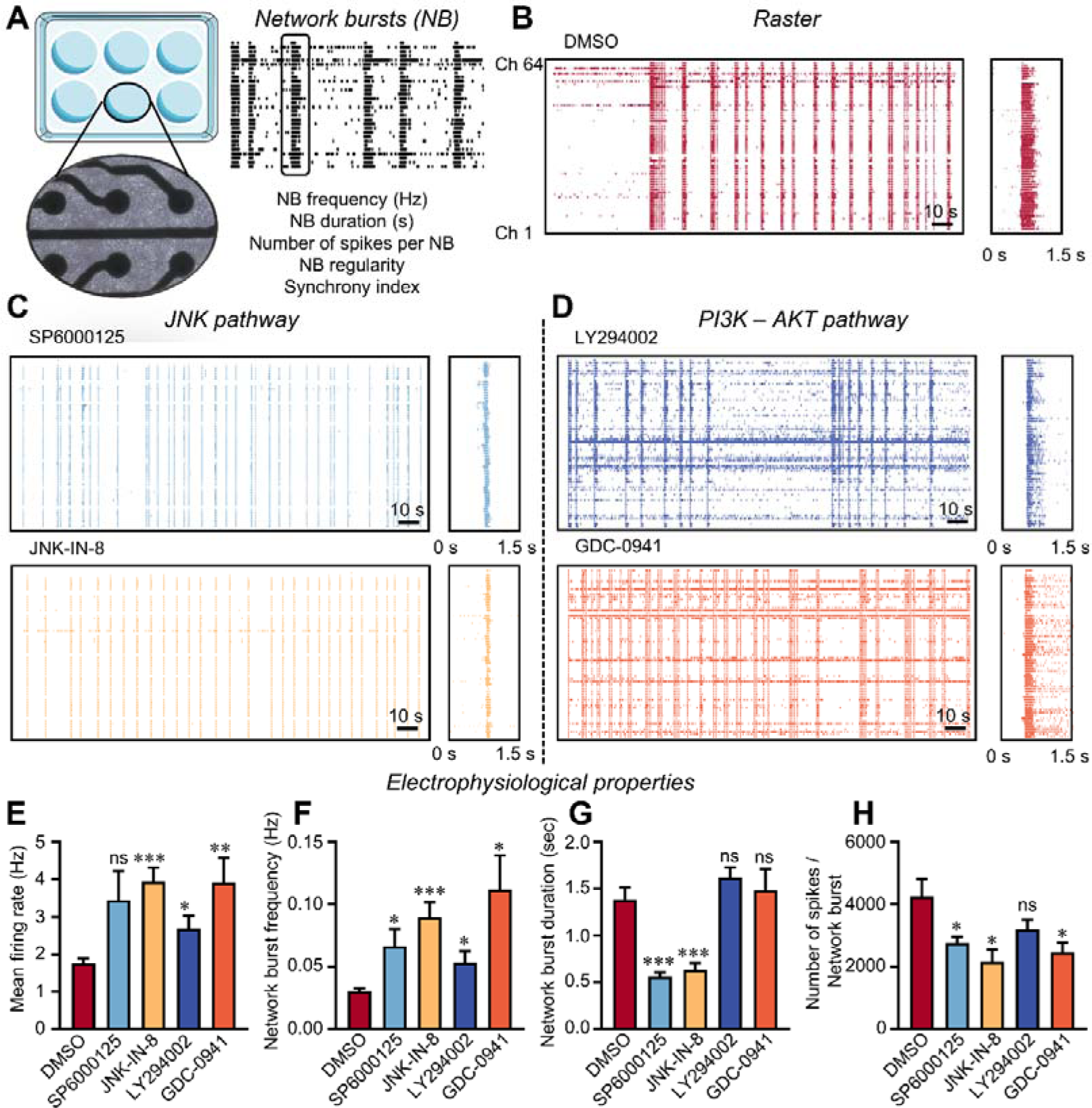
JNK inhibitors influence spontaneous electrophysiological activity of neuronal networks. **A.** Schematic representation of neuronal networks cultured on 6-well MEA. Schematic representation of spontaneous activity patterns measured on MEA. **B.** Representative raster plots showing 200 s of activity from DMSO culture at DIV 21. Right box shows representative network burst. **C.** Representative raster plots of JNK inhibitors for 200s recorded by MEA at DIV 21. Right box shows representative network burst of JNK inhibitors. **D.** Representative raster plots showing 200 s of electrophysiological activity recorded from PI3K-AKT inhibitors cultures at DIV 21. **E-H.** The (E) mean firing rate (MFR), (F) network burst frequency (NBF), (G) network burst duration (NBD) and (H) number of spikes per network burst were quantified. Recording time: 10 min. Electrophysiological data from 4 independent batches. *n*=10 for DMSO, *n*=9 for SP6000125, *n*=9 for JNK-in-8. *n*=8 for LY294002. *n*=10 for GDC-0941. All numerical data are means ± SD. Numbers in bars list the number of independent experiments analyzed. Statistical significance was examined by Student’s *t* test (\**P*<0.05, \*\**P*<0.01, \*\*\**P*<0.001).

Electrophysiological data was continuously recorded from DIV 14 to DIV 49. During neurodevelopment, the MFR of JNK inhibitors SP600025 and JNK-IN-8 began to decrease after reaching the peak at 21 days and showed a significant decrease compared with the DMSO starting from the fifth week (Figure 4D, *P*<0.05). Similarly, the NBF showed a similar trend to MFR after inhibiting JNK pathway (Figure 4A-E). Additionally, the NBD of JNK inhibitors SP600025 and JNK-IN-8 presented a prominent decrease from the third to the fifth week (Figure 4F, *P*<0.05). Other network parameters did not markedly differ from the DMSO (Figure 4G-I, *P*>0.05). The JNK signaling pathway memorably contributes to the development of synchronized bursting within the neuronal network. Finally, the inhibitors LY294002 and GDC-0941 exhibited no effect on MFR of neuronal network development (Figure 4A-D). Furthermore, inhibitors LY294002 and GDC-0941 had higher NBF at 2-4 weeks (Figure 4E, *P*<0.05), but their difference in NBD were not statistically significant (Figure 4F, *P*>0.05). Inhibitors LY294002 and GDC-0941 showed a prominent aggrandizement in NBD at 4-7 weeks (Figure 4F, *P*<0.05). Unexpectedly, the inhibition of the PI3K signaling pathway observably reduced the synchrony of neuronal networks after 5-7 weeks (Figure 5I, *P*<0.05). The emphasis of the PI3K pathway in network development and maintenance of network synchrony was further verified.

**Figure 4.**
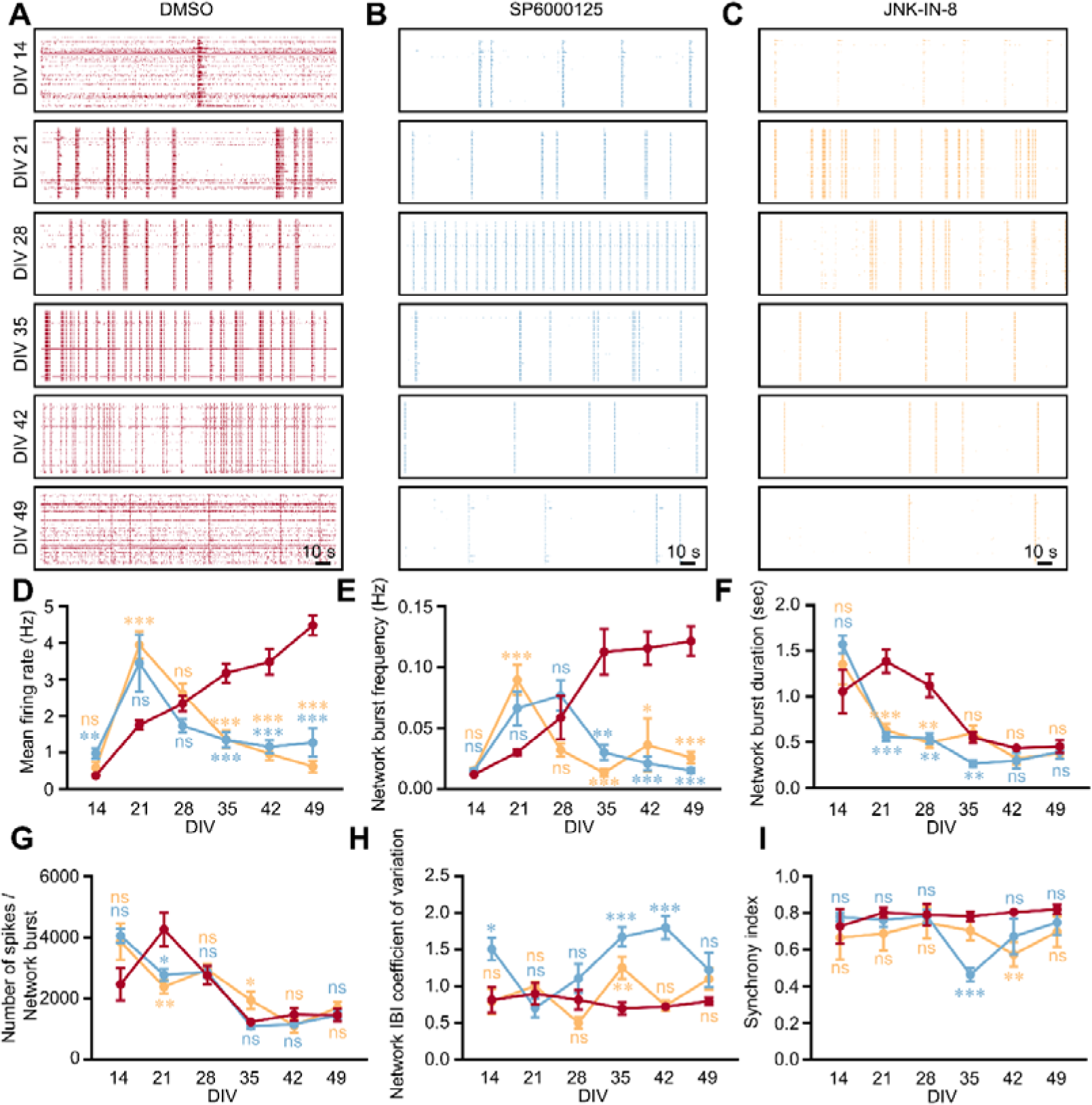
Development of spontaneous network activity in JNK inhibitors cultures. **A.** Representative raster plots of DMSO for 200 s recorded by MEA from DIV 14 to DIV 49. **B.** Representative raster plots of SP6000125 for 200s recorded by MEA from DIV 14 to DIV 49. **C.** Representative raster plots of JNK-IN-8 for 200s recorded by MEA from DIV 14 to DIV 49. **D-I.** Quantification of network parameters as indicated. Recording time: 10 min. Electrophysiological data from 4 independent batches. *n*=10 for DMSO, *n*=9 for SP6000125, *n*=9 for JNK-IN-8. All numerical data are means ± SD. Numbers in bars list the number of independent experiments analyzed. Statistical significance was examined by Student’s *t* test (\**P*<0.05, \*\**P*<0.01, \*\*\**P*<0.001).

**Figure 5.**
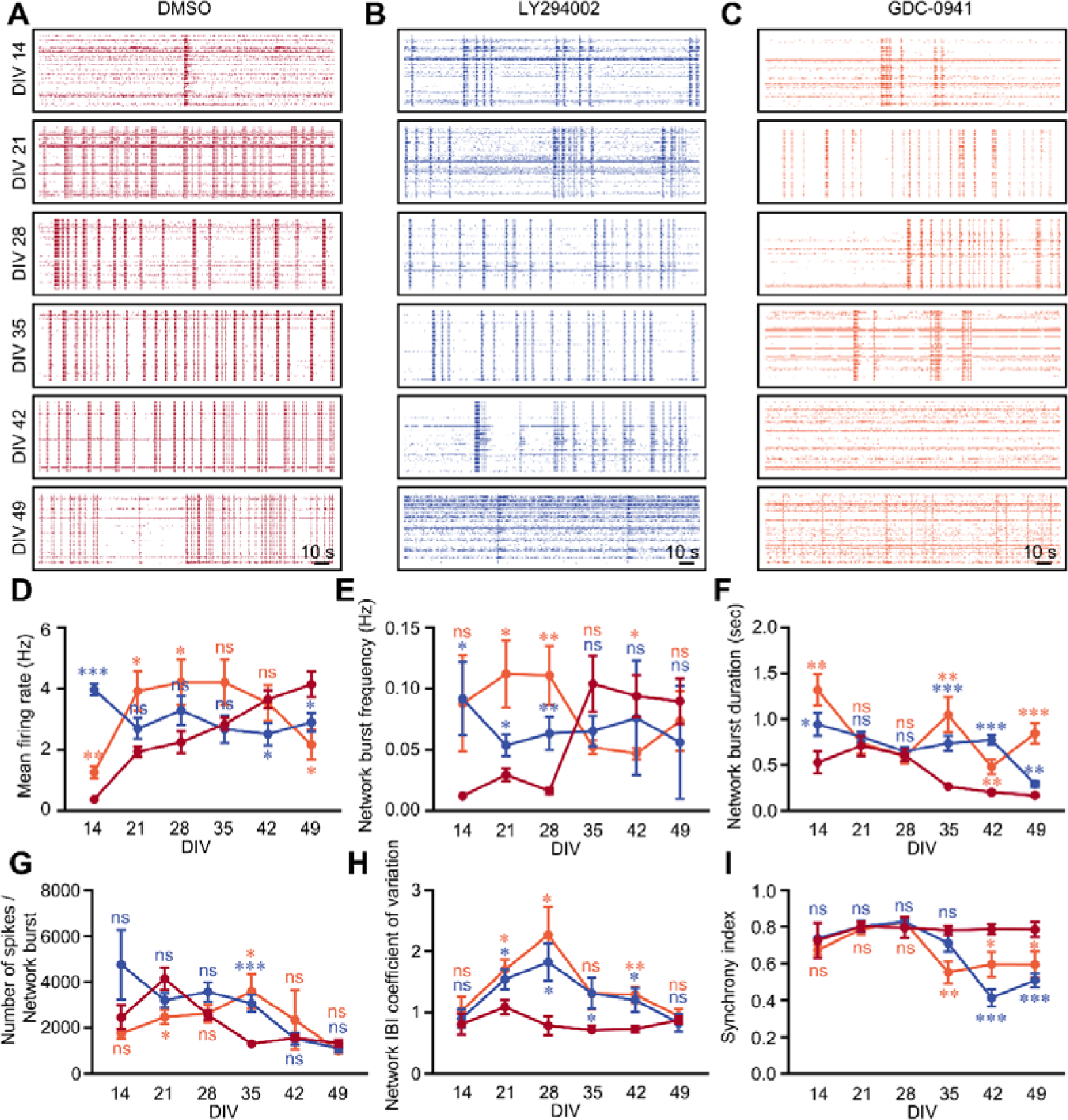
Development of spontaneous network activity in PI3K-AKT inhibitors cultures. **A.** Representative raster plots of DMSO for 200 s recorded by MEA from DIV 14 to DIV 49. **B.** Representative raster plots of LY294002 for 200 s recorded by MEA from DIV 14 to DIV 49. **C.** Representative raster plots of GDC-0941 for 200 s recorded by MEA from DIV 14 to DIV 49. **D-I:** Quantification of network parameters as indicated. Recording time: 10 min. Electrophysiological data from 4 independent batches. *n*=10 for DMSO, *n*=8 for LY294002. *n*=10 for GDC-0941. All numerical data are means ± SD. Numbers in bars list the number of independent experiments analyzed. Statistical significance was examined by Student’s *t* test (\**P*<0.05, \*\**P*<0.01, \*\*\**P*<0.001).

### Inhibition of JNK and PI3K pathways altered transcriptomic profile

Molecular changes in neuronal network phenotypic alterations resulting from inhibition of the JNK and PI3K-AKT pathways were also explored. RNA sequencing (RNA-seq) was conducted at DIV 15. Differentially expressed genes (DEGs) [FDR < 0.05, |log2FC| > log2 (2)] were detected in all treatment networks via DESeq2, as compared to dimethyl sulfoxide (DMSO, as control, Figure S4A). The transcriptional profiles’ principal component analysis (PCA) showed distinct biological replicate clustering for each treatment (Figure S4B). Quantification of DEGs display that a series of genes were down-/up-regulated in treated cultures compared to the control. DEGs identified in all treatments were remarkably comparable and mostly correlated to axon dendrite and synapse according to Gene Ontology (GO) annotation evaluated for cellular components (CC) (Figure 6A-B). GO annotation of DEGs identified from inhibitors SP6000125 and JNK-IN-8 networks exhibited substantial similarity in their homologous biological processes (BPs), particularly during system development (Figure 6C-D). GSEA-GO analysis showed that SP6000125 was in connection with axons and myelination, synaptic transmission, synaptic vesicle exocytosis and postsynaptic membrane (Figure S5A). However, the main function of JNK-IN-8 in neurons primarily involves mitochondrial BPs (Figure S5B). Thence, altered network development might be driven predominantly by the JNK pathway. Also, the BPs associated with inhibitors LY294002 and GDC-0941 were similar, mainly about axon ensheathment and system development (Figure 6E-F). Detected DEGs were related to alike molecular functions (MF), including ion binding and protein binding (Figure 6E-F). However, GO annotation assessed for CC suggested that the PI3K-AKT pathway was connected with axon and myelin sheath (Figure 6E-F). GSEA-GO analysis showed that the PI3K-AKT pathway was relevant to voltage-gated ion channel activity and neuronal synaptic transmission (Figure S6). Therefore, PI3K-AKT pathway commanded the neuronal structures and function.

**Figure 6.**
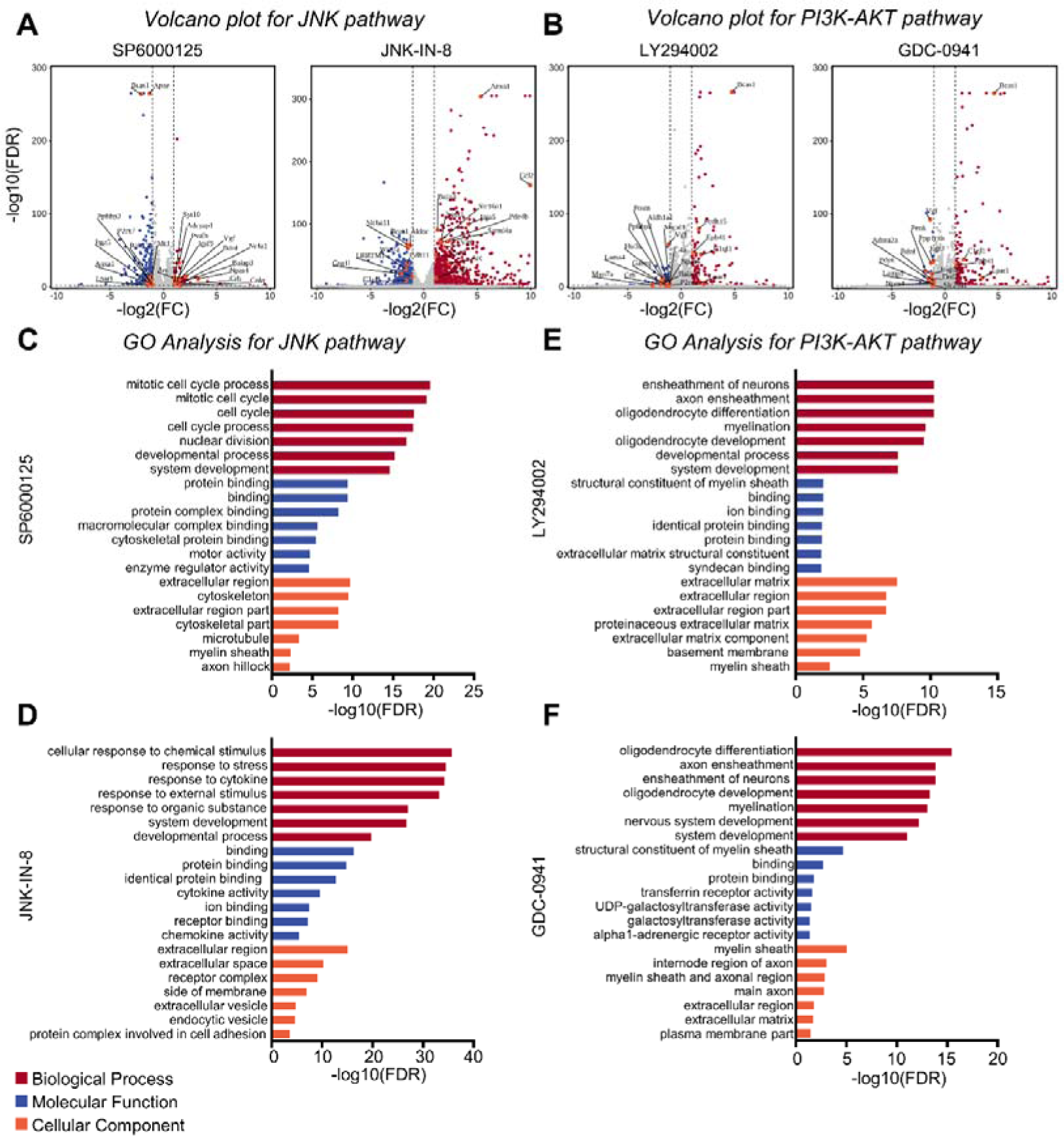
Multiple signaling pathways inhibitors cause deregulation of genes involved in neuronal and synaptic function and development. **A.** Volcano plots showing differentially expressed genes (DEGs) in JNK pathway inhibitors. Differential expression is defined by FDR<0.05 and abs(log_2_FC) > 0.3. Downregulation genes is blue points. Upregulation genes is red points. The orange points are labeled with the gene name associated with synapse. **B.** Volcano plots showing DEGs in PI3K-AKT pathway inhibitors. **C.** Gene Ontology (GO) term analysis of SP6000125 treatment associated with DEGs. **D.** GO term analysis of JNK-IN-8 treatment associated with DEGs. **E.** GO term analysis of LY294002 treatment associated with DEGs. **F.** GO term analysis of gdc-0941 treatment associated with DEGs. 3 replicates for each treatment.

What’s more, 251 genes shared commonalities among different JNK inhibitors (Figure S7A). Remarkably, these 251 genes demonstrated an enrichment of GO terms associated with myelination and axon function (Figure S7C). 150 genes associated with PI3K pathway were also identified (Figure S7B), and they correlative were development, axon and myelination (Figure S7D). Consequently, pathway related genes represented parallel functions in harmonizing neuronal structures and activity, notably enrich genes, directly influencing neuronal and synaptic function.

### Inhibition of JNK and PI3K pathways declined the expression of postsynaptic membrane receptors

Excitatory and inhibitory receptors are important in the spontaneous network activity and functional maturation of neuronal. Hence, on the 14th day, the excitatory receptors alpha-amino-3-hydroxy-5-methyl-4-isoxazolepropionic acid receptor (AMPAR) and N-methyl-D-aspartate receptor (NMDAR) were identified, along with the protein expression of the inhibitory receptors gamma-aminobutyric acid type A receptors (GABAARs). The JNK pathway inhibitor JNK-IN-8 significantly inhibited the protein expression of excitatory receptors AMPAR and NMDAR1 (Figure 7A, *P*<0.05). Similarly, the JNK pathway inhibitor SP6000125 also dramatically suppressed the expression of AMPAR and NMDAR1 (Figure 7B, *P*<0.01). Surprisingly, S6000125 also prominently inhibited the excitatory receptor GABAAR (Figure 7B, *P*<0.01). Inhibiting PI3K pathway with GDC-0941 and LY294002 led to conspicuous impairment in the excitatory receptor GABAAR (Figure 7C-D, *P*<0.05), but exhibited no arresting impact on inhibitory receptor proteins. Consequently, JNK and PI3K pathways exert an influence not only on synapse formation but also in post-synaptic membrane receptor types.

**Figure 7.**
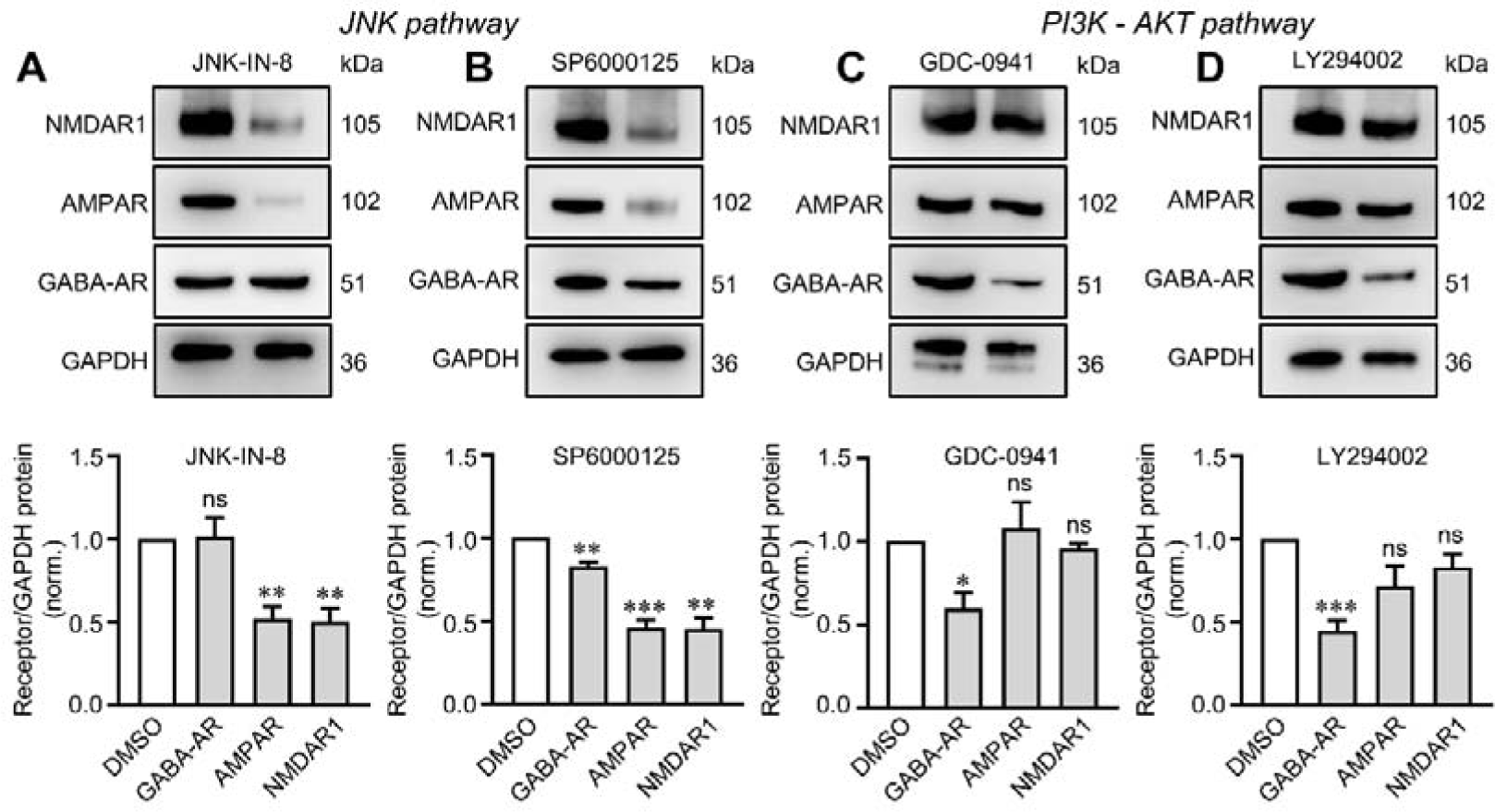
Multiple signaling pathways influence the expression of postsynaptic membrane receptors. Top, representative immunoblots images; below, quantifications. **A.** Immunoblotting was performed to assess GABA-AR, AMPAR, and NDAR1 expression levels following treatment with the JNK inhibitor JNK-IN-8, respectively revealing detected bands at 51 kDa, 102 kDa and 105 kDa. **B.** Immunoblotting was performed to assess GABA-AR, AMPAR, and NDAR1 expression levels following treatment with the JNK inhibitor SP6000125, severally revealing detected bands at 51 kDa, 102 kDa and 105 kDa. **C.** Immunoblotting was performed to assess GABA-AR, AMPAR, and NDAR1 expression levels following treatment with the PI3K inhibitor GDC-0941, respectively revealing detected bands at 51 kDa, 102 kDa and 105 kDa. **D.** Immunoblotting was performed to assess GABA-AR, AMPAR, and NDAR1 expression levels following treatment with the PI3K inhibitor LY294002, severally revealing detected bands at 51 kDa, 102 kDa and 105 kDa. All numerical data are means ± SD. Numbers in bars list the number of independent experiments analyzed. Statistical significance was examined by Student’s *t* test (\**P*<0.05, \*\**P*<0.01, \*\*\**P*<0.001, *n*=3 independent experiments).

## Discussion

Our previous research has focused on the identification and characterization of multiple signaling pathways mediating synapse formation by HSF assay (*4*). Although most synaptic adhesion molecules induce pre- or postsynaptic specializations in the HSF assay, many of them are not essential for synapse formation *in vivo* (*18*). However, they actually contribute to organizing specific synaptic properties, highlighting notable limitations of this assay. Any transsynaptic signal activated by synaptic adhesion molecules can instigate the organization of synapses (*19*). Most importantly, the synapses formed in non-neuronal cells, such as HEK293T, are not “real” synapses in HSF assays (*20*). It seems likely that PKA, JNK and PI3K pathways are involved in promoting synapse formation, but their involvement between genuine neurons has not yet been explored.

In this study, pharmacological methods and MEA were employed to investigate the involvement of canonical signal transduction pathways in synapse formation within cultured neurons. During the maturation of cultured neuronal networks, the period spanning DIV7-14 emerges as a pivotal phase of synaptogenesis which is a intricate process governing synapse formation (*21*). Therefore, pathway inhibitors were added during DIV7-14 to inhibit the critical process of synapse formation. In cultured neurons, the formation of pre-/post-synaptic specialization necessitates PKA and JNK pathway, two canonical protein kinase pathways, while the PI3K pathway is merely specifically required for post-synaptic specialization. However, inhibiting the PKA signal through pharmacological methods significantly affected neuronal morphology, suggesting that PKA pathway not only modulates synapse formation at DIV 7-14, but also disrupted fundamental processes like neuronal growth and development (Figure 1 and S2). Given the multifaceted involvement of the PKA pathway in various BPs, its effect on synapse formation during the DIV7-14 was not highlight in this survey. Molecular intervention may be a more efficient and targeted method for inhibition of PKA pathway. Therefore, based on the limitations of pharmacological interventions where a train of disrupters was used to confirm the specificity of pathways, this study focused on the role of JNK and PI3K signal in synapse formation. Although the PI3K pathway did not affect presynaptic puncta density, it significantly affected the size of presynaptic puncta. Thus, PI3K pathway is also involved in presynaptic specialization in cultured neurons (Figure 2). On account of the interconnection of these pathways and their cross-talk at various levels, they might possess aggregation to facilitate synaptogenesis processes. It is still of great heuristic value to explore the specific action sites of various signaling pathways in synaptic formation. In addition, two inhibitors were used for each signal pathway in order to obtain more stable and convincing results. However, due to the unique pharmacological characteristics of each pathway inhibitor, strict consistent results were not obtained in the detection of some synapse-related proteins, but similar trends were obtained.

MEA tools provide valuable insights into neuronal network phenotypes (*22*). The relationship between network plasticity and spontaneous NBs in cultured neuronal networks have been identified in our antecedent studies (*23*). MEA technology enables robust monitoring of extracellular network activities, uniformly sampled over several months, which fabricates opportunities for precise pharmacological manipulations combined with long-term activity recordings *in vivo* (*24*). Spontaneous network activity laid the infrastructure of functional networks during early central nervous system development (*17*). Spontaneous NBs dynamics triggered by the unique modifications of each pathway inhibitor have been observed in cultured neural networks. The network phenotypes after inhibiting JNK and PI3K pathways at DIV 21 showed striking similarities, resulting in a higher MFR and NBF compared with DMSO, or hyperactivity. Generally, the hyperactivity of neural networks can be attributed to: (i) variation in excitatory/inhibitory (E/I) balance resulting from synaptic signaling between neurons, and (ii) transformation in intrinsic electrophysiological properties of neurons within the networks (*25*). The formation of E/I synapses is regulated by various signaling pathways. The E/I imbalance of synapses extremely enticed abnormal neural network activities. MFR and NBs of neuronal networks were increased after GABAA receptor inhibition (*10*), while the duration of spontaneous NBs was reduced with NMDAR blockade (*11*), which aligns closely with our experiment. Consequently, JNK pathway obstructs postsynaptic GABAA receptors and NMDA receptors, while the PI3K signal disturbs GABAA receptors.

Multidose chronic blockers have greater flexibility in manipulating synaptic receptor activation than single dose acute treatment, enabling progressive adjustments in the E/I balance (*11*). The developmental trajectory of JNK pathway shows that impeding JNK signal significantly reduced MFR and NBF of mature networks, broke firing regularity, but maintained their synchronization. A similar increase in NBs similarity was previously proposed, effectively reducing the number of burst initiation zones in postnatal cortical cultures by restraining protein kinase C (PKC) (*26*). PKC is recognized for enhancing AMPAR conductance through the phosphorylation of AMPAR subunits (*27*). The modulations of integral network activity after PKC inhibition may be synchronized with JNK blocking, thus further adjusting AMPAR function. Besides, functional inhibition of GABAA receptors cannot maintain the hyperactivity of neuronal network for a long time. After the network matures, impaired GABAA receptor function affected the developmental trajectory of network. The *in vitro* dynamics of NBs may be impact by diversified cellular mechanisms and phenomena, including synaptic scaling (*28*), AMPAR/NMDAR trafficking extra-synaptic receptors (*29*) and even ephaptic coupling (*30*). After PI3K signaling suppression, either NBD or synchrony index of the network varied memorably after 5 weeks of network development. The activity patterns, pertinence and synchrony of neurons in the network are principally regulated by NMDA receptors, while NBD is controllable by GABAA receptors. Therefore, PI3K signal affects the later development of the network through excitatory and inhibitory postsynaptic receptors. Certainly, changes in network firing activity are related to E/I ratio of synapses, indicating that E/I balance profoundly interferences network dynamics. NBs activity is mediated by different pathways. First, the JNK pathway prevailingly mediates the neuronal network’s MFR, NBF and NBD. Second, NBs lasts longer and synchronicity lessens when PI3K pathway is damaged.

Transcriptomics systematically reveals the transcriptional landscape of JNK and PI3K signal. As expected, JNK pathway is mainly involved in BPs including growth and development on account of enrichment in synaptic, cytoskeletal and axonal elements in cortical neurons. Meanwhile, PI3K signal primarily participated in myelination and neurodevelopment processes. Furthermore, PI3K signaling restriction significantly affected voltage-gated ion channel activity, synaptic transmission and synaptic plasticity of neurons. Changes in the electrophysiological activity of neuronal networks after PI3K inhibition may be closely related to the alteration of these BPs (*31*). PI3K pathway regulates GABAergic receptors related to NBD, resulting in phenotypic defect of the electrophysiology. An intriguing finding is that 251 overlapping genes related to the JNK signaling pathway and 150 overlapping genes related to the PI3K signaling pathway were identified. The main functions of these genes are to target neurodevelopmental processes and myelination. Myelination plays a crucial role in the nervous system by protecting neurons (*32*), increasing the speed of nerve impulse conduction (*33*), preventing interference from other electrical impulses, and maintaining the strength of nerve impulses (*34*). Computational models show that neuronal activity-dependent changes in myelination promote neural synchronization (*35*), which is responsible for the reduced synchrony in the later stages of neural network development after inhibiting the PI3K signaling pathway. Additionally, the association of JNK and PI3K pathways with the cytoskeleton suggests their potential contribution to synapse formation through cytoskeletal regulation.

## Conclusions

In conclusion, JNK and PI3K pathways involved in synapse formation influence the electrophysiology of neuronal networks. Firstly, canonical JNK pathway is necessary for pre- and post-synaptic specialization, while the PI3K has a profound impact on postsynaptic specialization and pre-synaptic puncta size. Secondly, JNK signal commands MFR, NBF, NBD and the development of network, while inhibiting the PI3K gives rise to a longer burst duration and disadvantaged network synchrony. Thirdly, deficiency in JNK and PI3K pathways contributes to the deregulation of genes controlling neuronal and synaptic processes. Finally, JNK pathway primarily influences the expression of excitatory receptors, whereas PI3K signal is associated with inhibitory receptors.

## Supplementary information

**Additional file: Figure S1:** Minimal inhibitory concentration gradient screening. **Figure S2:** PKA inhibitors H89 and PKI impair the formation of pre- and post-synaptic side in primary cultures of cortical neurons. **Figure S3:** Quantification of the network IBI coefficient of variation and synchrony index at DIV 21. **Figure S4:** KEGG analysis different inhibitors target anticipatory signal pathway. **Figure S5:** GSEA analysis of all genes about JNK pathway. **Figure S6:** GSEA analysis of all genes about PI3K-AKT pathway. **Figure S7:** Multiple signaling pathways overlap DEGs involved in neuronal function and development.

## Acknowledgments

Our gratitude extends to the Bioimaging Core of Shenzhen Bay Laboratory for their invaluable imaging support. Special appreciation is due to Bioimaging Core engineers Mei Yu and Shixian Huang for their assistance with the laser scanning confocal microscopy LSM 980. Additionally, we acknowledge Hailin Lu and Jiying Hu from the Drug Discovery Core of Shenzhen Bay Laboratory for their support with the MEA recording.

## Author Contributions

J.X.L and J.X designed experiment. J.X.L and W.S.H preformed experiment. J.X.L and Z.Q.Y wrote the manuscript. J.X.L and Z.Z.H analyzed the data. All authors contributed to this paper and approved the submitted version.

## Availability of data and materials

The datasets supporting the conclusions of this article are available from the author, without undue reservation.

## Declarations

### Ethical Approval and Consent to participate

The experimental animals were carried out in accordance with the guidelines issued by the Animal Ethics Committee of ShenZhen Bay Laboratory (Approval No: AEJX202201).

### Consent for publication

Not applicable.

## Reference

1. T. C. Südhof, Towards an Understanding of Synapse Formation. Neuron 100, 276–293 (2018).

2. T. C. Südhof, The cell biology of synapse formation. Journal of Cell Biology 220, (2021).

3. T. Biederer, P. Scheiffele, Mixed-culture assays for analyzing neuronal synapse formation. Nature Protocols 2, 670–676 (2007).

4. X. Jiang, R. Sando, T. C. Südhof, Multiple signaling pathways are essential for synapse formation induced by synaptic adhesion molecules. Proc. Natl. Acad. Sci. U. S. A. 118, (2021).

5. T. Biederer et al., SynCAM, a synaptic adhesion molecule that drives synapse assembly. Science 297, 1525–1531 (2002).

6. M. Frega et al., Neuronal network dysfunction in a model for Kleefstra syndrome mediated by enhanced NMDAR signaling. Nature Communications 10, (2019).

7. D. J. Bakkum et al., Tracking axonal action potential propagation on a high-density microelectrode array across hundreds of sites. Nature Communications 4, (2013).

8. P. C. Antonello et al., Self-organization of in vitro neuronal assemblies drives to complex network topology. Elife 11, (2022).

9. Y. L. Li et al., Characterization of synchronized bursts in cultured hippocampal neuronal networks with learning training on microelectrode arrays. Biosensors & Bioelectronics 22, 2976–2982 (2007).

10. Y. L. Li, W. Zhou, X. N. Li, S. Q. Zeng, Q. M. Luo, Dynamics of learning in cultured neuronal networks with antagonists of glutamate receptors. Biophysical Journal 93, 4151–4158 (2007).

11. H. Teppola, J. Acimovic, M. L. Linne, Unique Features of Network Bursts Emerge From the Complex Interplay of Excitatory and Inhibitory Receptors in Rat Neocortical Networks. Frontiers in Cellular Neuroscience 13, (2019).

12. I. Dias et al., Consolidation of memory traces in cultured cortical networks requires low cholinergic tone, synchronized activity and high network excitability. Journal of Neural Engineering 18, (2021).

13. A. Awasthi et al., Synaptotagmin-3 drives AMPA receptor endocytosis, depression of synapse strength, and forgetting. Science 363, 44-+ (2019).

14. W. D. Hale, T. C. Suedhof, R. L. Huganir, Engineered adhesion molecules drive synapse organization br. Proc. Natl. Acad. Sci. U. S. A. 120, (2023).

15. K. J. Gan, T. C. Südhof, SPARCL1 Promotes Excitatory But Not Inhibitory Synapse Formation and Function Independent of Neurexins and Neuroligins. Journal of Neuroscience 40, 8088–8102 (2020).

16. X. Jiang et al., A Small Molecule That Protects the Integrity of the Electron Transfer Chain Blocks the Mitochondrial Apoptotic Pathway. Molecular Cell 63, 229–239 (2016).

17. W. Plumbly, N. Patikas, S. F. Field, S. Foskolou, E. Metzakopian, Derivation of nociceptive sensory neurons from hiPSCs with early patterning and temporally controlled *NEUROG2* overexpression. Cell Reports Methods 2, (2022).

18. L. Chen, C. Nam, Postsynaptic assembly induced by neurexin-neuroligin interaction and transmitter release. Molecular Biology of the Cell 15, 96A–96A (2004).

19. T. C. Südhof, Molecular Neuroscience in the 21st Century: A Personal Perspective. Neuron 96, 536–541 (2017).

20. S. J. Lee et al., Presynaptic Neuronal Pentraxin Receptor Organizes Excitatory and Inhibitory Synapses. Journal of Neuroscience 37, 1062–1080 (2017).

21. N. E. Ziv, S. J. Smith, Evidence for a role of dendritic filopodia in synaptogenesis and spine formation. Neuron 17, 91–102 (1996).

22. E. Parnell et al., Excitatory Dysfunction Drives Network and Calcium Handling Deficits in 16p11.2 Duplication Schizophrenia Induced Pluripotent Stem Cell-Derived Neurons. Biological Psychiatry 94, 153–163 (2023).

23. X. L. Jia et al., Learning populations with hubs govern the initiation and propagation of spontaneous bursts in neuronal networks after learning. Frontiers in Neuroscience 16, (2022).

24. D. Pré et al., Development of a platform to investigate long-term potentiation in human iPSC-derived neuronal networks. Stem Cell Reports 17, 2141–2155 (2022).

25. S. Wang et al., Loss-of-function variants in the schizophrenia risk gene *SETD1A* alter neuronal network activity in human neurons through the cAMP/PKA pathway. Cell Reports 39, (2022).

26. S. Okujeni, U. Egert, Self-organization of modular network architecture by activity-dependent neuronal migration and outgrowth. Elife 8, (2019).

27. M. A. Jenkins, S. F. Traynelis, PKC phosphorylates GluA1-Ser831 to enhance AMPA receptor conductance. Channels 6, 60–64 (2012).

28. M. F. Fong, J. P. Newman, S. M. Potter, P. Wenner, Upward synaptic scaling is dependent on neurotransmission rather than spiking. Nature Communications 6, (2015).

29. R. C. Carroll, R. S. Zukin, NMDA-receptor trafficking and targeting: implications for synaptic transmission and plasticity. Trends in Neurosciences 25, 571–577 (2002).

30. M. Martinez-Banaclocha, Ephaptic Coupling of Cortical Neurons: Possible Contribution of Astroglial Magnetic Fields? Neuroscience 370, 37–45 (2018).

31. B. K. Unda et al., Impaired OTUD7A-dependent Ankyrin regulation mediates neuronal dysfunction in mouse and human models of the 15q13.3 microdeletion syndrome. Molecular Psychiatry 28, 1747–1769 (2023).

32. P. H. H. Lopez et al., Myelin-associated glycoprotein protects neurons from excitotoxicity. Journal of Neurochemistry 116, 900–908 (2011).

33. S. Ravera, A. M. Morelli, I. Panfoli, Myelination increases chemical energy support to the axon without modifying the basic physicochemical mechanism of nerve conduction. Neurochemistry International 141, (2020).

34. T. W. Chapman, R. A. Hill, Myelin plasticity in adulthood and aging. Neuroscience Letters 715, (2020).

35. G. Bonetto, D. Belin, R. T. Káradóttir, Myelin: A gatekeeper of activity-dependent circuit plasticity? Science 374, 838-+ (2021).

