## Supplemental Figure for "JNK and PI3K signaling pathways mediate synapse formation and network spontaneous activities in primary neurons"

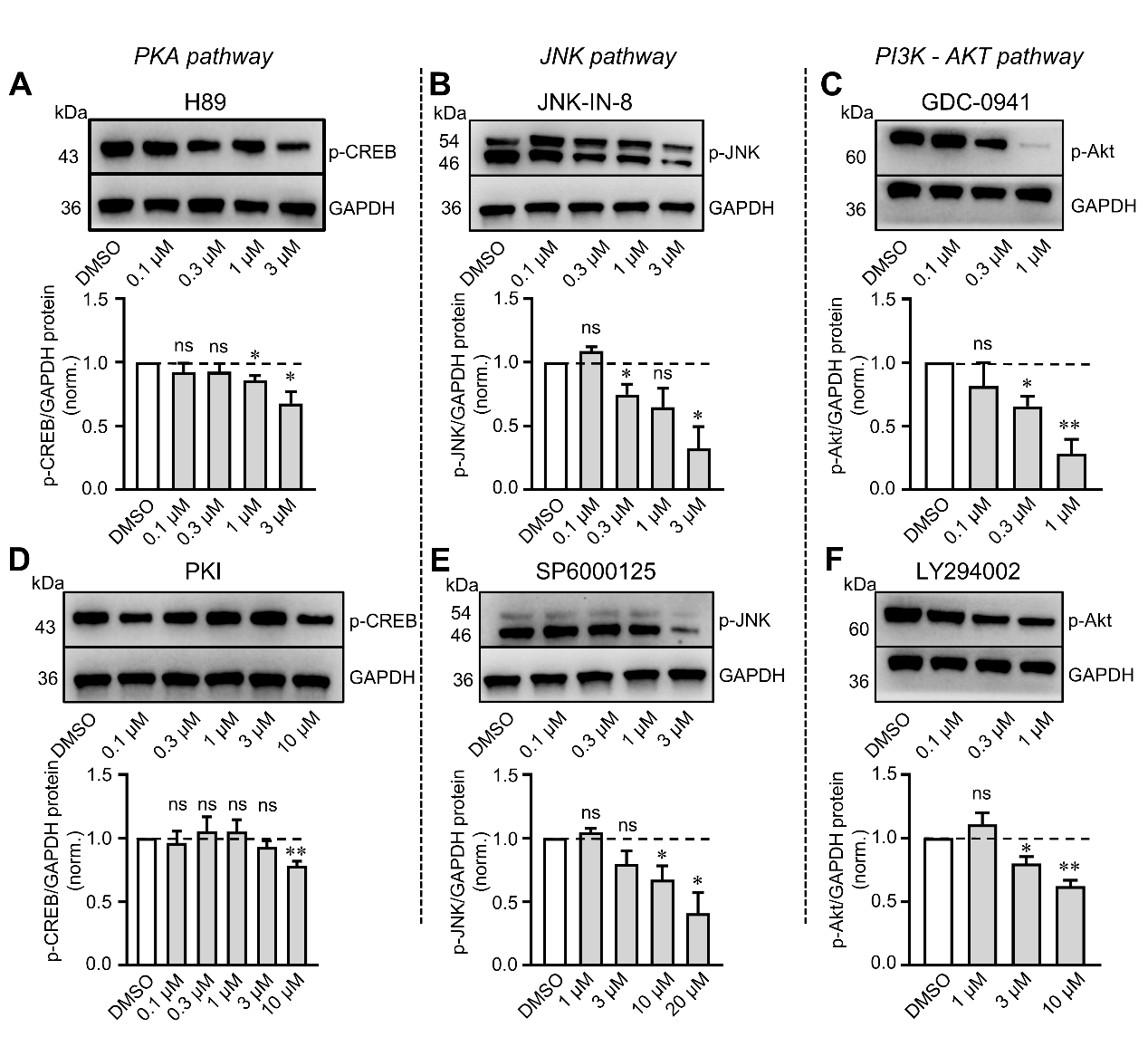

**Supplement Figure 1.** Minimal inhibitory concentration gradient screening.

Top, representative immunoblots images; below, quantifications. **A.** Immunoblotting for p-CREB to screen minimal inhibitory concentration of PKA inhibitor H89 was assayed by western blot with detected bands at 43 kDa. **B.** Immunoblotting for p-JNK to screen minimal inhibitory concentration of JNK inhibitor JNK-IN-8 was assayed by western blot with detected bands from 46 kDa to 54 kDa. **C.** Immunoblotting for p-AKT to screen minimal inhibitory concentration of PI3K inhibitor GDC-0941 was assayed by western blot with detected bands at 60 kDa. **D.** Immunoblotting for p-CREB to screen minimal inhibitory concentration of PKA inhibitor PKI was assayed by western blot with detected bands at 43 kDa. **E.** Immunoblotting for p-JNK to screen minimal inhibitory concentration of JNK inhibitor SP6000125 was assayed by western blot with detected bands from 46 kDa to 54 kDa. **F.** Immunoblotting for p-AKT to screen minimal inhibitory concentration of PI3K inhibitor LY294002 was assayed by western blot with detected bands at 60 kDa. Summary graphs of protein levels normalized for GAPDH as an internal standard. All numerical data are means ± SD. Numbers in bars list the number of independent experiments analyzed. Statistical significance was examined by Student’s t test (**P*<0.05, ***P*<0.01, ****P*<0.001, *n*=3 independent experiments).

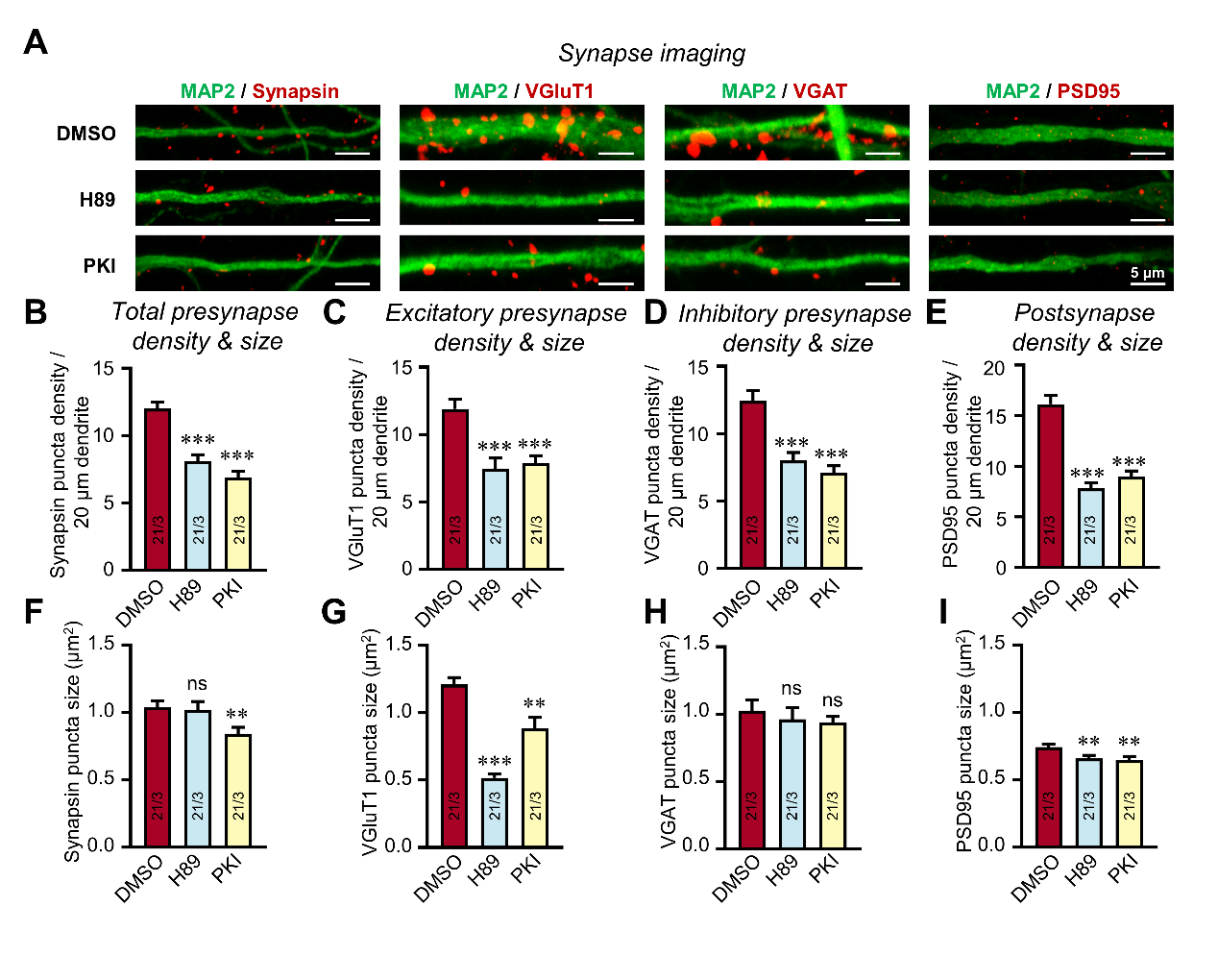

**Supplement Figure 2.** PKA inhibitors H89 and PKI impair the formation of pre- and post-synaptic side in primary cultures of cortical neurons.

**A**. Representative images showing PKA inhibitors treatment decrease both pre- and post-synapse density. Neurons were immunostaining for Synapsin1/2 (synapse marker), VGluT1 (excitatory presynaptic marker), VGAT (inhibitory presynaptic marker), PSD95 (postsynaptic marker) and MAP2 (dendritic marker) at DIV15. **B, F**. Quantifications showing that PKA inhibitors H89 and PKI decrease total pre-synapse density and size. **C, G**. Quantifications showing that PKA inhibitors decrease excitatory pre-synapse density and size. **D, H**. Quantifications showing that PKA inhibitors decrease inhibitory pre-synapse density but does not alter inhibitory pre-synaspse size. **E, I**. Quantifications showing that PKA inhibitors has significant effect on post-synapse density and size. All numerical data are means ± SD. Numbers in bars list the number of independent experiments/cells analyzed. Statistical significance was examined by Student’s t test (**P*<0.05, ***P*<0.01, ****P*<0.001).

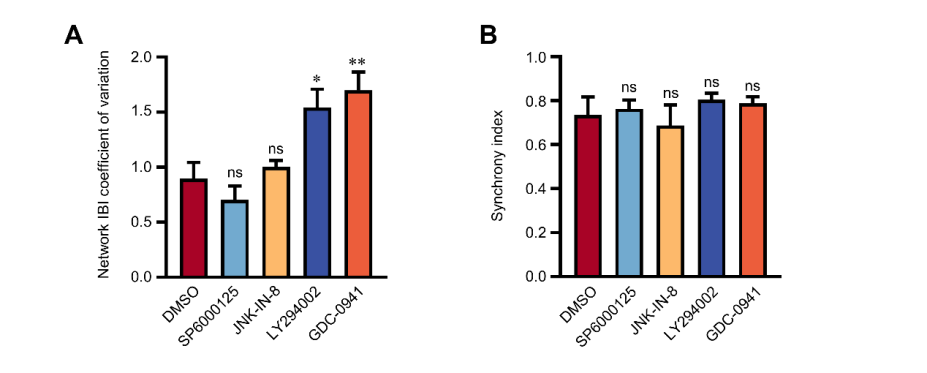

**Supplement Figure 3.** Quantification of the network IBI coefficient of variation and synchrony index at DIV 21.

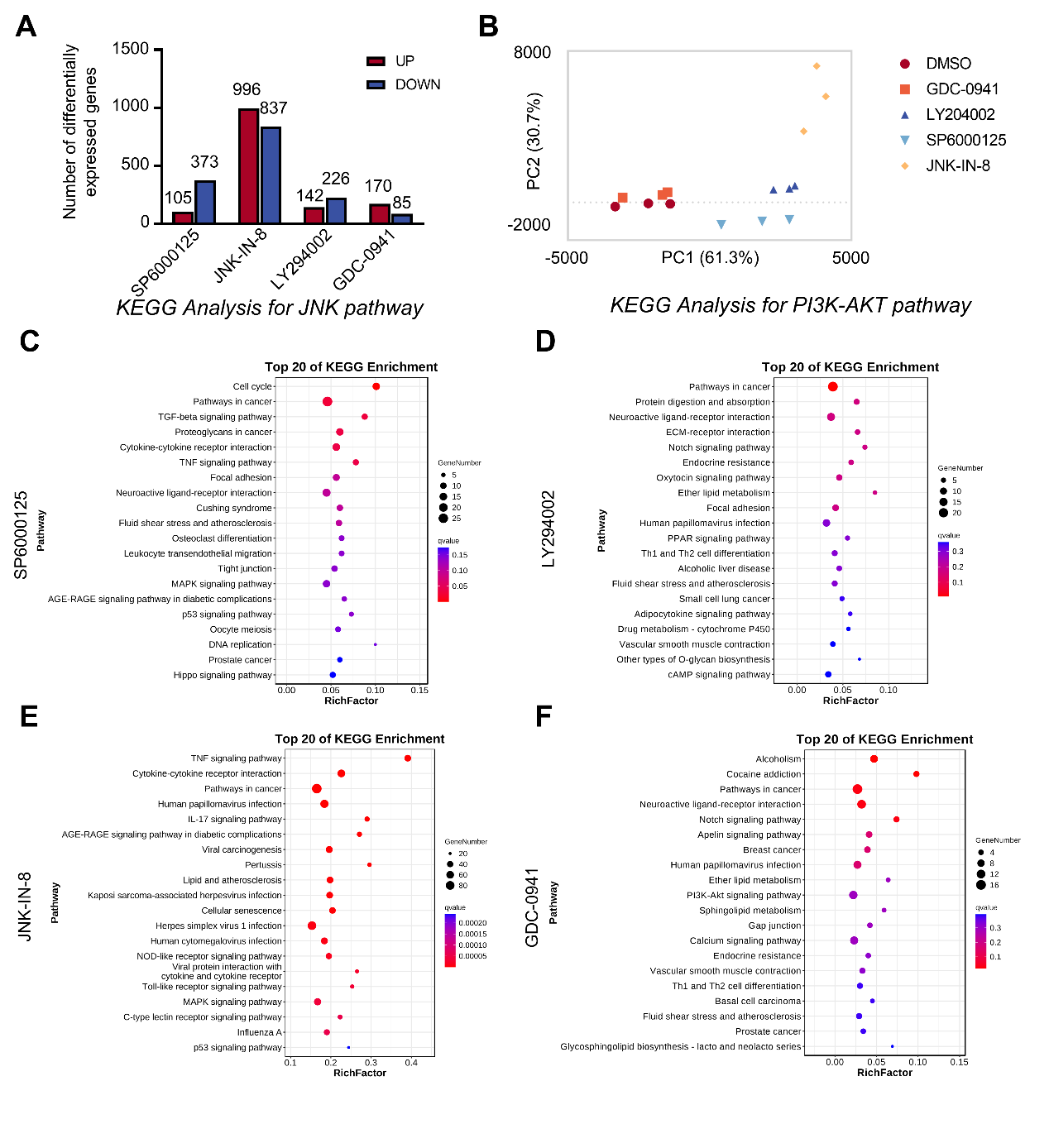

**Supplement Figure 4.** KEGG analysis different inhibitors target anticipatory signal pathway.

**A.** Number of differentially expressed (DE) genes in the different conditions, as compared to DMSO. **B.** PCA showing tight clustering of 3 replicates for each treatment. **C.** KEGG analysis of DEGs associated with SP6000125 treatment. **D.** KEGG analysis of DEGs associated with LY294002 treatment. **E.** KEGG analysis of DEGs associated with JNK-IN-8 treatment. **F.** KEGG analysis of DEGs associated with GDC-0941 treatment.

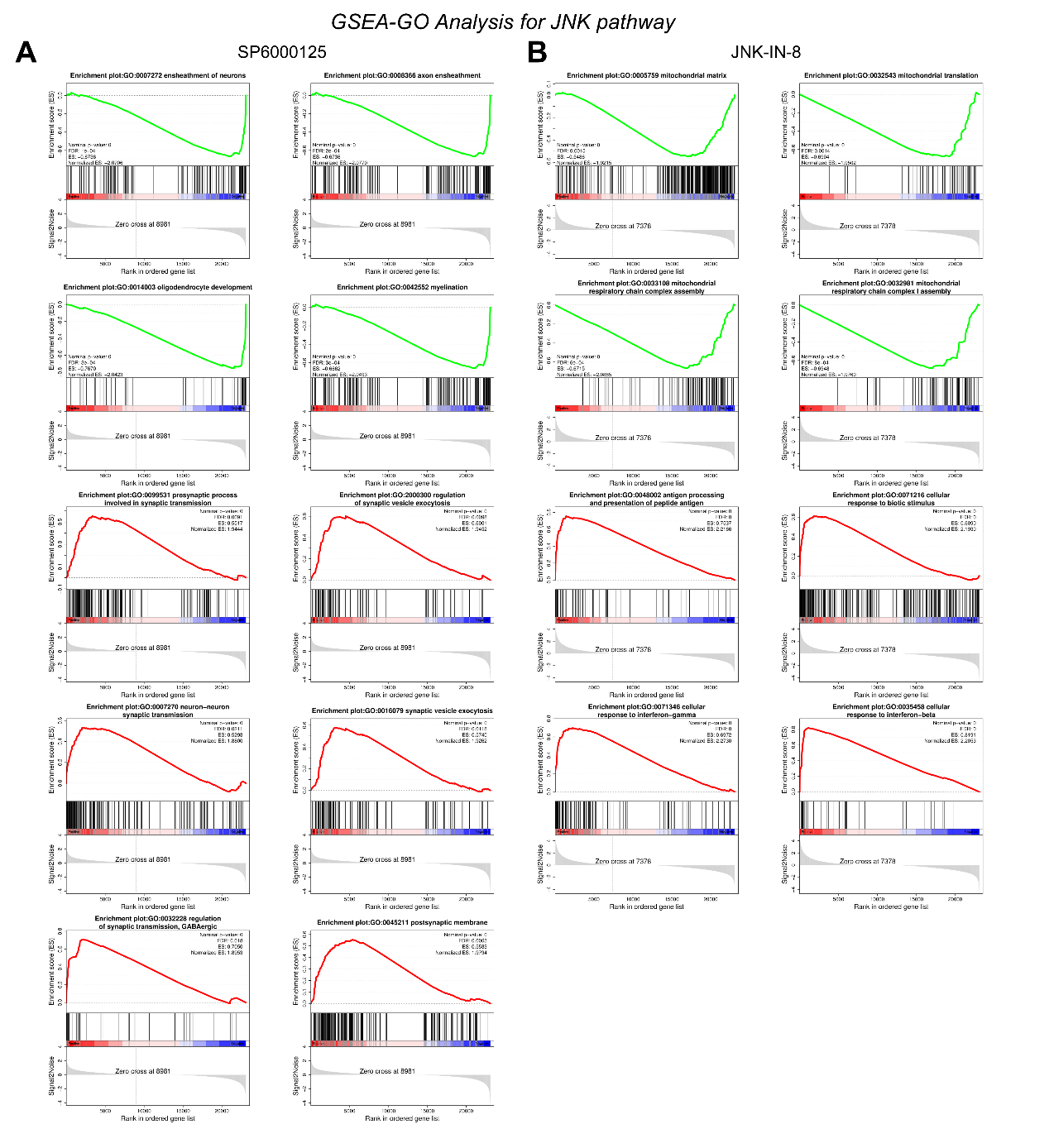

**Supplement Figure 5.** GSEA analysis of all genes about JNK pathway

**A.** SP6000125 mainly involves pathways related to neuronal function and synaptic transmission. **B.** JNK-IN-8 mainly involves mitochondrial related functions.

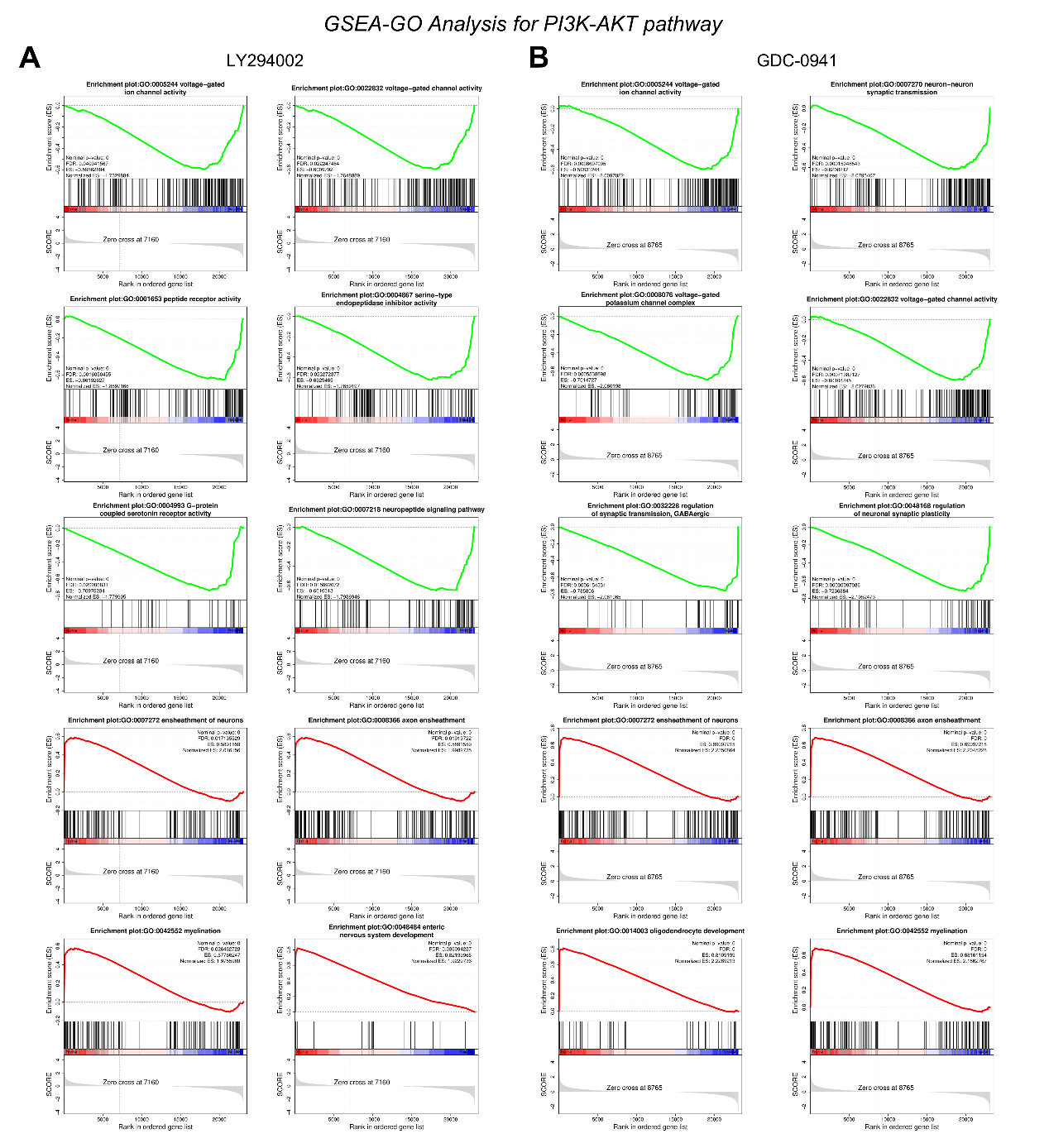

**Supplement Figure 6.** GSEA analysis of all genes about PI3K-AKT pathway

**A.** LY294002 treatment mainly involves neuronal functions and development. **B.** GDC-0941 treatment mainly associated with neuron and synaptic function and myelination.

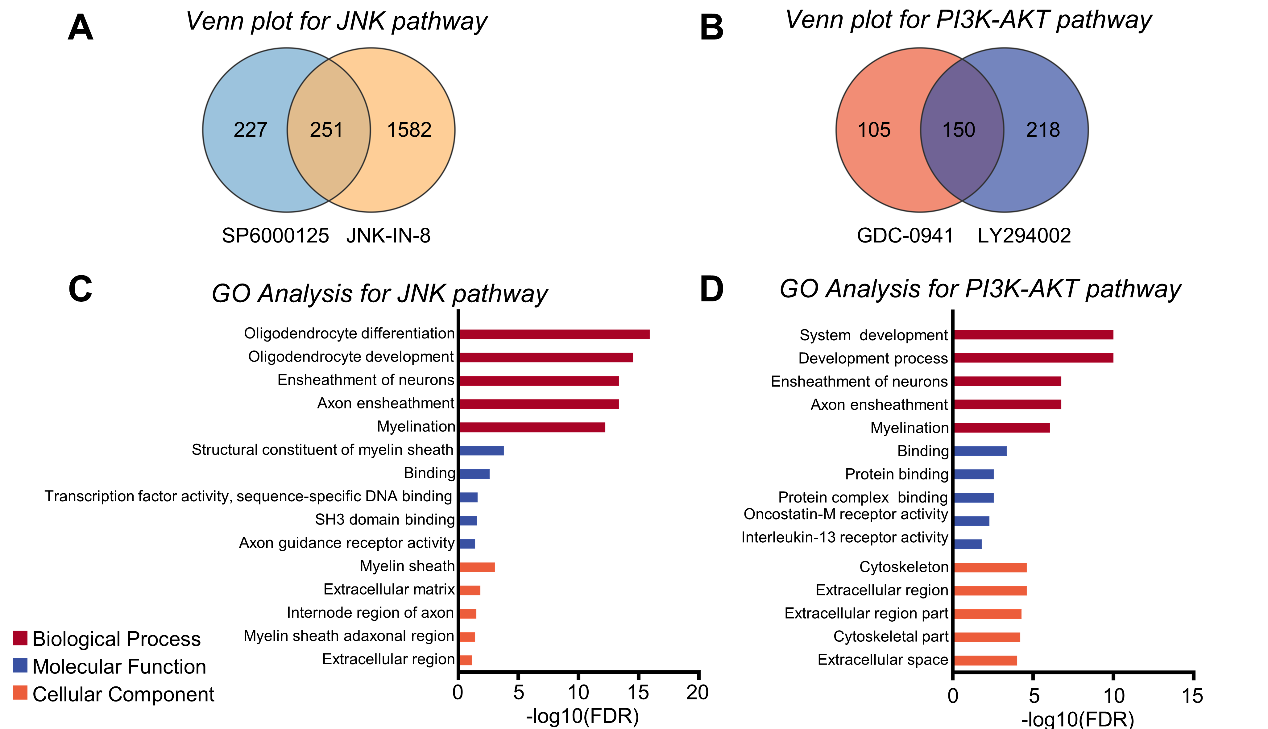

**Figure S7.** Multiple signaling pathways overlap DEGs involved in neuronal function and development.

**A.** JNK pathway inhibitors SP6000125 and JNK-IN-8 overlap DEGs is 251. **B.** PI3K-AKT pathway inhibitors GDC-0941 and LY294002 overlap DEGs is 150. **C.** Gene Ontology (GO) term analysis of 251 DEGs. **D.** GO term analysis of 150 DEGs.
